# Replication gaps underlie BRCA-deficiency and therapy response

**DOI:** 10.1101/781955

**Authors:** Nicholas J. Panzarino, John Krais, Min Peng, Michelle Mosqueda, Sumeet Nayak, Samuel Bond, Jennifer Calvo, Ke Cong, Mihir Doshi, Matt Bere, Jianhong Ou, Bin Deng, Lihua Julie Zhu, Neil Johnson, Sharon B. Cantor

## Abstract

Cancers that are deficient in BRCA1 or BRCA2 are hypersensitive to genotoxic agents, including platinums and other first-line chemotherapeutics. The established models propose that these cancers are hypersensitive because the chemotherapies block or degrade DNA replication forks and thereby create DNA double strand breaks, both of which require functional BRCA proteins to prevent or resolve by mechanisms termed fork protection (FP) or homologous recombination (HR). However, recent findings challenge this dogma because genotoxic agents do not initially cause DNA double strand breaks or stall replication forks. Here, we propose a new model for genotoxic chemotherapy in which ssDNA replication gaps underlie the hypersensitivity of BRCA deficient cancer, and we propose that defects in HR or FP do not. Specifically, we observed that ssDNA gaps develop in BRCA deficient cells because DNA replication is not effectively restrained in response to genotoxic stress. Moreover, we observe gap suppression (GS) by either restored fork restraint or by gap filling, both of which conferred resistance to therapy in tissue culture and BRCA patient tumors. In contrast, restored HR and FP were not sufficient to prevent hypersensitivity if ssDNA gaps were not eliminated. Together, these data suggest that ssDNA replication gaps underlie the BRCA cancer phenotype, “BRCAness,” and we propose are fundamental to the mechanism of action of genotoxic chemotherapies.

## Introduction

Mutations in the hereditary breast cancer genes, BRCA1 and BRCA2, first demonstrated that cancer is a genetic disease in which susceptibility to cancer could be inherited (King et al., 1986). In addition to breast cancer, mutated BRCA1 or BRCA2 also cause a predisposition to other cancer types, including breast, ovarian, pancreatic, and colorectal cancers. Importantly, cancers with mutated BRCA genes are hypersensitive to cisplatin, a first-line anti-cancer chemotherapy that has been the standard of care for ovarian cancer for over 40 years (Munnell, 1968). BRCA deficient cancers are thought to be hypersensitive to cisplatin due to their inability to repair cisplatin-induced DNA double strand breaks (DSBs) by homologous recombination (HR) (Li and Heyer, 2008). Accordingly, it is proposed that the DNA breaks are created when replication forks collide with the cisplatin-DNA crosslinks, causing the fork to collapse into DSBs (Feng and Jasin, 2017). This broken-fork-model was further supported by reports that mutations in the BRCA genes also lead to defective fork protection (FP), which are thought to render forks vulnerable to fork collapse and subsequent DSB induction (Schlacher et al., 2011; Schlacher et al., 2012). Correspondingly, chemoresistance in BRCA cancer is proposed to occur when either HR or FP is restored, with the latter largely preventing DSBs and therefore eliminating the requirement for HR.

However, recent findings challenge the fundamental premise that DSBs are the critical lesion for cisplatin sensitivity. Notably, cisplatin does not initially cause replication forks to stall and collapse (Huang et al., 2013; Mutreja et al., 2018). Moreover, recent findings indicate that cisplatin toxicity in triple negative breast cancer is unrelated to loss of DNA repair factors (Heijink et al., 2019). In addition, in several distinct models restored FP fails to restore cisplatin resistance, suggesting FP is uncoupled from the mechanism of resistance (Feng and Jasin, 2017) (Cantor and Calvo, 2018). Most saliently, indicating that the underlying sensitizing lesion may in fact not be a DSB, HR proficient cells show cisplatin hypersensitivity (Wang et al., 2015). Moreover, in addition to cisplatin, BRCA deficient cells and patient tumors have been found to be hypersensitive to a wide range of genotoxic agents that were previously thought to be distinct, including doxorubicin, PARPi, and others agents that do not generate DSBs such as oxaliplatin (Bruno et al., 2017). Taken together, these findings indicate an opportunity to revise the current framework for both BRCAness as well as the mechanism of action of multiple first-line chemotherapies.

Here, we present evidence that BRCAness derives from single stranded DNA (ssDNA) formation, and not from the failure to repair or prevent the induction of DNA double strand breaks due to defects in HR or FP. Specifically, we observed in BRCA deficient cells that ssDNA gaps develop because DNA replication is not effectively restrained in response to genotoxic stress. Moreover, we observed gaps could be suppressed by either restored fork restraint or by gap filling, both of which conferred resistance to therapy in tissue culture and BRCA patient tumors. In contrast, restored FP or HR were not sufficient to prevent hypersensitivity if ssDNA gaps were not eliminated. Together, these data indicate that ssDNA replication gaps underlie the BRCA cancer phenotype, “BRCAness,” and we propose are fundamental to the mechanism of action of genotoxic chemotherapies.

## Results

To analyze the mechanism underlying the hypersensitivity of BRCA deficient cancers to chemotherapy, we monitored the immediate response of DNA replication to stress using DNA fiber assays. In DNA fiber assays, cells are exposed to the nucleotide analogs IdU and CldU, which are incorporated into nascent DNA as the cells replicate, and are subsequently detected by immunofluorescence when the DNA is spread on a slide. Accordingly, the dynamics of replication can be measured by quantifying the length of the DNA that contains analog labels under different conditions. Here, we measured the lengths of the labeled DNA when the cells were exposed to hydroxyurea (HU), which depletes the nucleotide pool by inhibiting the protein ribonucleotide reductase, an enzyme that is critical for the proper synthesis of DNA nucleotides. Importantly, the replication stress is induced by low dose HU (0.5 mM) without fully depleting nucleotide pools (Koc et al., 2004), and compared to treatment with other genotoxic chemotherapies (such as DNA crosslinkers) produces higher quality DNA fibers. Thus, we exposed BRCA proficient and deficient cancer cells to HU and analog at the same time in order to compare the immediate response to replication stress with and without the BRCA genes. As shown in **Figure 1A**, the parental PEO1 cancer cell line has a truncated BRCA2 protein and is hypersensitive to cisplatin, whereas the BRCA2 proficient PEO1 reversion cell line, C4-2, is resistant to cisplatin (Sakai et al., 2009). The cell lines were incubated with 5-lodo-2’-deoxyuridine (IdU) for 30 minutes as a control to label regions of active replication, followed by 5-chloro-2’-deoxyuridine (CldU) for 2h in the presence of HU to monitor the immediate response of DNA replication to genotoxic stress. We observed longer CldU tracks in the BRCA2-deficient PEO1 cells compared to the BRCA2-proficient C4-2 cells, indicating replication in PEO1 cells had failed to fully slow during HU (**Figure 1B**), and suggests that BRCA2 is required to effectively restrain replication during stress. Indeed, further suggesting that replication restraint is a function of the BRCA proteins, we observed that a BRCA2 deficient Chinese hamster cell line (VC-8) (Schlacher et al., 2011), and two BRCA1 deficient breast cancer lines (HCC1937 and MDA-MB-436) showed that DNA replication was arrested more effectively during stress when cells were complemented with the respective wild type BRCA genes (**Figure S1.1A-C**). Similarly, as an additional control, we also depleted BRCA2 in BRCA2-proficient C4-2 cells, and again found that forks were not fully arrested after HU (**Figure S1.1D-F**). In addition, we also confirmed these results with an experiment that measures total analog fluorescence in the nucleus, rather than on individual DNA fibers, and therefore functions as a readout of global cellular replication. Indeed, in BRCA2 deficient cells, we observed an increase in EdU positive cells during replication stress compared to BRCA2 proficient cells, further supporting that BRCA2 is required to restrain replication during stress (**Figure 1C, S1C**). Lastly, we treated cells with cisplatin instead of HU, and again found that BRCA2-deficient PEO1 cells had longer replication tracts compared to BRCA2 proficient C4-2 (**Figure S1.1G**), indicating that restraint defects are not restricted to stress induced by HU.

**Figure 1:**
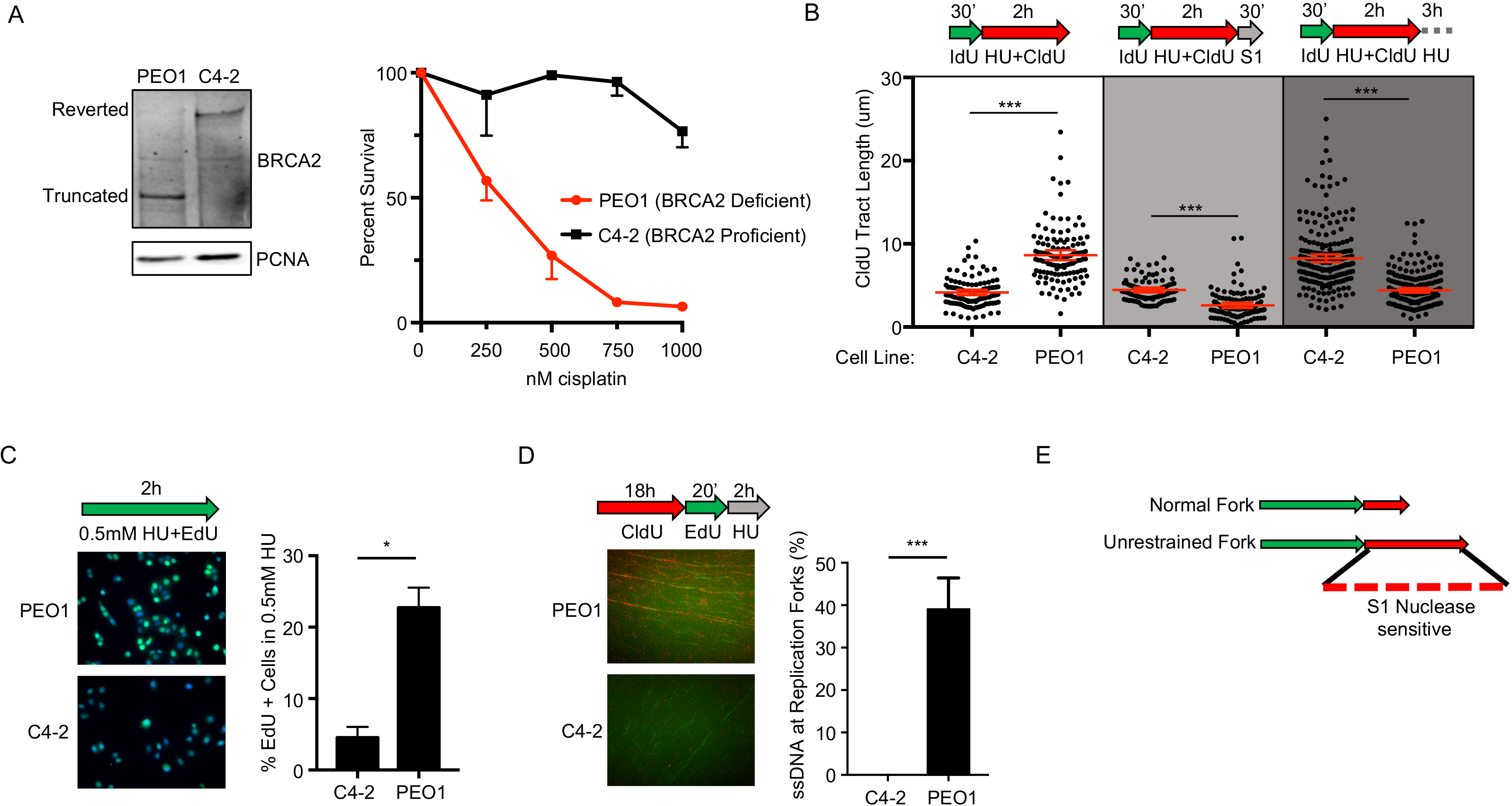
BRCA2 deficient cancer cells fail to restrain replication in the presence of stress, generating regions of ssDNA gaps that are destroyed after continued exposure. **A)** Left, Western blot detects truncated BRCA2 protein in BRCA2 deficient PEO1 cells, and detects full-length BRCA2 protein in BRCA2 proficient C4-2 cells that are derived from PEO1 cells. Right, cell survival assay confirms PEO1 cells are hypersensitive to cisplatin compared to C4-2 cells. **B)** Schematic and quantification of CIdU track length shows (white panel) that PEO1 cells fail to arrest replication in the presence of stress. These regions are degraded by S1 nuclease (light grey panel), and are also destroyed after continued exposure to replication stress (dark grey panel). Each dot represents one fiber. Experiments were performed in biological triplicate with at least 100 fibers per replicate. Statistical analysis according to two-tailed Mann-Whitney test; *** p < 0.001. Mean and 95% confidence intervals are shown. **C)** Schematic, representative images and quantification of nuclear imaging identifies a greater percentage of EdU positive cells in PEO1 as compared to C4-2. p < 0.05 (*) as determined by t-test of biological triplicate experiments. **D)** Nondenaturing fiber assay identifies exposed ssDNA in newly replicating regions after stress in PEO1, but not C4-2 cells. Regions of active replication were detected with EdU-ClickIT (green signal). p < 0.01 (***) as determined by t-test of biological triplicate experiments. **E)** Model of fiber assay interpretation.

Given that failure to restrain replication is associated with genome wide ssDNA gaps (Hashimoto et al., 2010; Henry-Mowatt et al., 2003; Kolinjivadi et al., 2017; Zellweger et al., 2015), we hypothesized that failure to fully restrain replication in response to stress in BRCA deficient cells would result in poorly replicated regions that contain ssDNA. To test this hypothesis, we performed the DNA fiber assay followed by incubation with S1 nuclease. S1 cuts at ssDNA regions and secondary DNA structures as an indicator of poor-quality DNA (Quinet et al., 2017). Indeed, labelled nascent DNA tracks were S1 sensitive in BRCA2-deficient PEO1 cells, but not in the BRCA2-proficient C4-2 cells (**Figure 1B**). These S1 sensitive nascent DNA regions were also degraded after continued exposure to replication stress indicating that nascent DNA in regions behind the fork (>10kb) are degraded under continued stress. (**Figure 1B and E**). In addition, to directly confirm that these S1 sensitive regions in the PEO1 cells contained ssDNA, we employed a non-denaturing DNA fiber assay that detects ssDNA in regions of active DNA replication. Indeed, this assay confirmed that ssDNA (detected as CldU signal) was present adjacent to newly replicating EdU positive regions in the BRCA2 deficient PEO1 cells, but not in the BRCA2 proficient C4-2 cells (**Figure 1D**). Similarly, BRCA1 deficient cancer cells also displayed DNA replication tracks that were sensitive to S1 nuclease after treatment with HU (**Figure S1.1H**). Thus, BRCA deficient cancer cells fail to fully restrain replication in the presence of stress, creating ssDNA regions (**Figure 1E**) that are degraded after additional exposure to stress.

We further hypothesized that ssDNA gaps confer chemosensitivity in BRCA cancer, and that mechanisms of chemoresistance would lead to gap suppression (GS). Indeed, we found that gaps were suppressed when chemoresistance was achieved by depletion of the chromatin remodeling enzyme CHD4 in BRCA2 deficient PEO1 cells (**Figure 2A**) (Guillemette et al., 2015). Specifically, when CHD4 was depleted, we observed significantly suppressed S1 nuclease sensitivity compared to the control, and nascent DNA tracks were not degraded after continued exposure to HU (**Figure 2B**). In agreement, ssDNA (measured as CldU signal) adjacent to regions of active replication were significantly reduced when CHD4 was depleted (**Figure 2D**). We also observed these DNA fiber phenotypes with a second shRNA reagent to CHD4 (**Figure S2A,B, see Figure 3E**). However, when CHD4 was depleted, replication restraint in response to stress was not re-established. Instead, replication tracks were significantly longer in CHD4-depleted PEO1 cells, compared to PEO1 control or C4-2 cells (**Figure 2B**). Moreover, in agreement with increased replication track length, analysis of global cellular replication by EdU incorporation demonstrated that CHD4 depleted PEO1 cells also had a greater number of EdU positive cells after HU treatment as compared to control PEO1 or C4-2 cells. As an additional control, EdU positive cells were similar in unchallenged controls, suggesting the replication differences occur in response to stress (**Figure 2C, S2A-E**). Thus, gaps and chemosensitivity were suppressed by CHD4 depletion, but fork restraint was not restored, indicating replication was further mis-regulated (**Figure 2E**). Critically, in C4-2 cells the BRCA2 reversion mutation (Sakai et al., 2009) also suppresses ssDNA gaps (**Figure 1B, 2B**), suggesting that chemoresistance is achieved by either gap filling or restored fork slowing.

**Figure 2:**
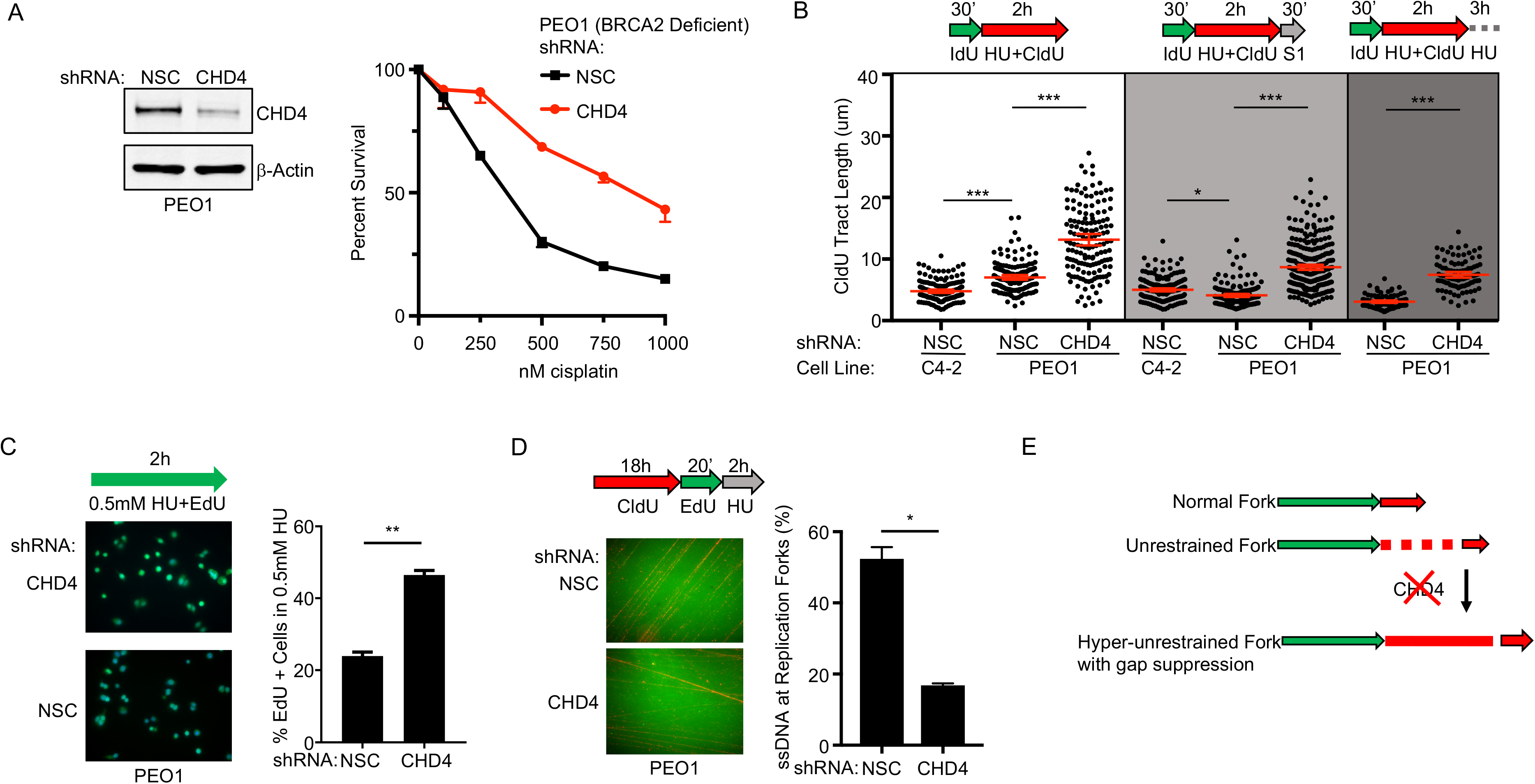
CHD4 depletion suppresses ssDNA gaps but does not restore fork restraint. **A)** Left, Western blot confirms CHD4 is depleted by shRNA compared to non-silencing control (NSC) in BRCA2 deficient PEO1. Right, cell survival assay confirms PEO1 with depleted CHD4 are resistant to cisplatin compared to PEO1 NSC. **B)** Schematic and quantification of CIdU track length shows (white panel) that PEO1 with depleted CHD4 increase replication in the presence of stress (white panel). These regions are protected from S1 nuclease (light grey panel), and are also protected after continued exposure to replication stress (dark grey panel). Each dot represents one fiber. Experiments were performed in biological triplicate with at least 100 fibers per replicate. Statistical analysis according to two-tailed Mann-Whitney test; *** p < 0.001. Mean and 95% confidence intervals are shown. **C)** Schematic, representative images and quantification of nuclear imaging identifies a greater percentage of EdU positive cells in CHD4 depleted PEO1 as compared to NSC. p < 0.01 (**) as determined by t-test of biological triplicate experiments. **D)** Nondenaturing fiber assay identifies that ssDNA in newly replicating regions after stress is reduced when CHD4 is depleted in PEO1 cells. Regions of active replication were detected with EdU-ClickIT (green signal). p < 0.05 (*) as determined by t-test of biological duplicate experiments. **E)** Model of fiber assay interpretation.

Our data indicate that suppression of ssDNA replication gaps in BRCA deficient cancer could confer chemoresistance. To address this possibility, we sought to identify additional genes similar to CHD4 that confer chemoresistance by GS. Thus, we performed quantitative mass spectrometry proteomics comparing the CHD4-interactome between BRCA2 deficient vs proficient cells after cisplatin treatment (**Figure 3A**). In BRCA2 deficient PEO1 cells, CHD4 interacted with several proteins known, when lost, to confer chemoresistance without restoring HR, including PARP1, EZH2, and FEN1 (**Figure 3B**) (Guillemette et al., 2015; Meghani et al., 2018; Rondinelli et al., 2017). We also identified the CHD4-interacting protein ZFHX3 (Chudnovsky et al., 2014), whose depletion we found enhanced cisplatin resistance in PEO1 cells (**Figure 3C**). Moreover, in BRCA2 deficient ovarian cancers, low mRNA levels of ZFHX3 predicted poor tumor-free survival (**Figure 3D**) as previously found for CHD4, EZH2, and FEN1 (Guillemette et al., 2015; Meghani et al., 2018; Rondinelli et al., 2017). Strikingly, we observed that depletion of ZFHX3 or FEN1, or EZH2 inhibition, increased replication in the presence of HU (**Figure 3E**), as found for CHD4 depletion. Moreover, in the S1 nuclease assay, inhibition of EZH2 or depletion of ZFHX3 or FEN1 also protected regions of nascent DNA from S1 nuclease degradation in BRCA2 deficient cells, similar to the depletion of CHD4 (**Figure 3E, Figure S2A,F**). Together, these findings suggest that loss of CHD4, EZH2, FEN1, and ZFHX3 suppresses ssDNA gaps to confer chemoresistance.

**Figure 3:**
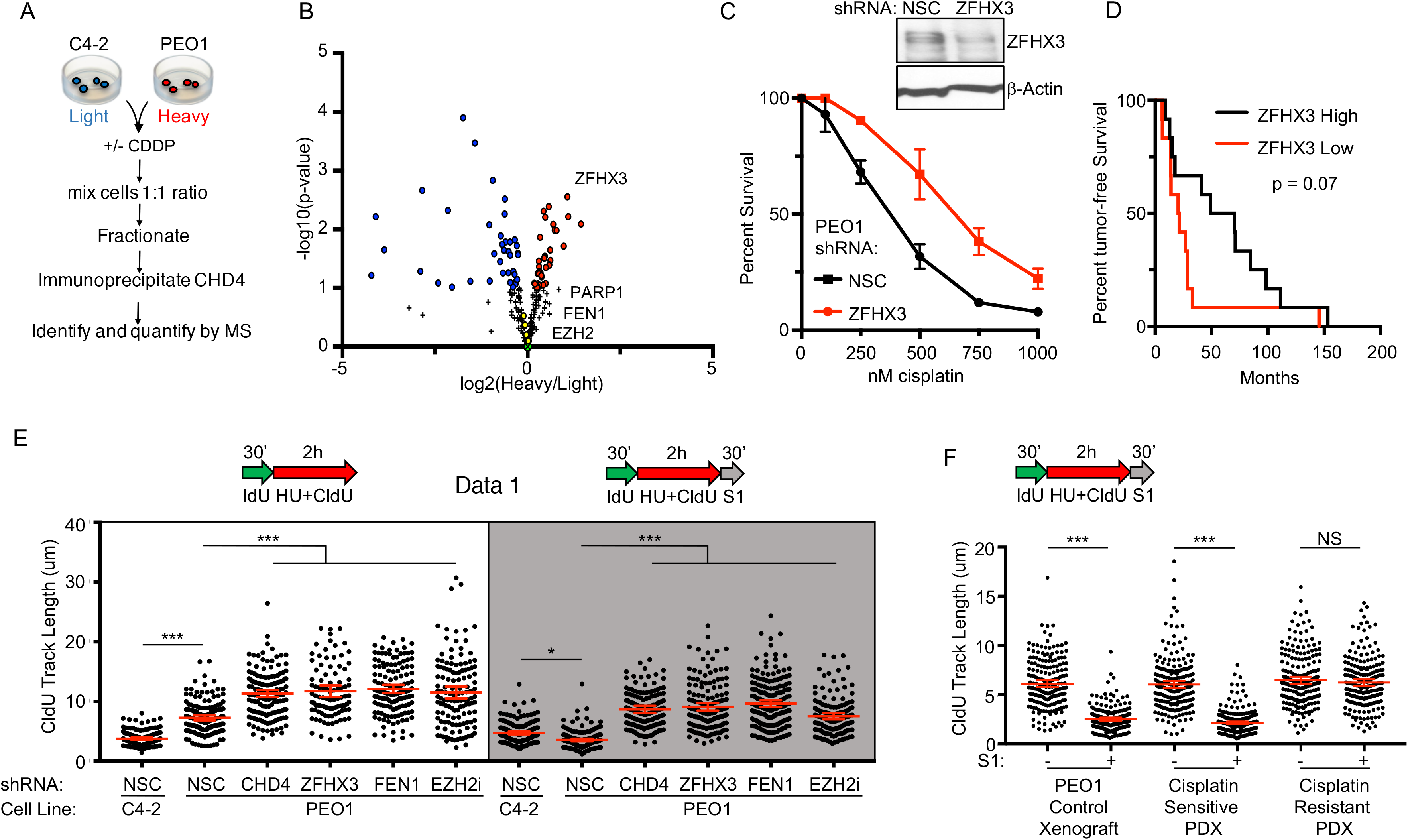
Suppression of ssDNA gaps accurately predicts poor therapy response in both cell culture and patient xenografts. **A)** Overview of the SILAC CHD4 immunoprecipitation experiment. **B)** SILAC immunoprecipitation reveals that CHD4 interacts with ZFHX3, FEN1, and EZH2 after cisplatin treatment. Red and blue circles are proteins significantly enriched in the CHD4 network of either PEO1 or C4-2 cells. Yellow circles are known CHD4 interacting partners (O’Shaughnessy and Hendrich, 2013). Black plus signs represent proteins not significantly enriched in the CHD4 network of either PEO1 or C4-2. Three biological replicates were performed; see methods section for statistical analysis. **C)** Western blot confirms ZFHX3 is depleted by shRNA in PEO1 as compared to NSC. Cell survival assay confirms PEO1 with depleted ZFHX3 are resistant to cisplatin compared to PEO1 NSC. **D)** Reduced ZFHX3 mRNA levels predict poor patient response to therapy (progression free survival) for BRCA2 deficient ovarian cancers from the TCGA database. **E)** Schematic and quantification of CldU track length shows that depletion of CHD4 (shRNA#61), ZFHX3 or FEN1, or inhibition of EZH2, increase replication in the presence of stress (white panel) and protect nascent DNA from S1 nuclease (gray panel). **F)** Schematic and quantification of CldU track length shows S1 fiber sensitivity is suppressed in BRCA1 deficient patient derived xenografts that have acquired chemoresistance. Each dot represents one fiber. Experiments were performed in biological triplicate with at least 100 fibers per replicate; the xenograft fiber assay was performed in duplicate. Statistical analysis according to twotailed Mann-Whitney test; *** p < 0.001. Mean and 95% confidence intervals are shown.

Next, we tested if ssDNA gaps could predict chemosensitivity and resistance in BRCA patient tumor samples. We utilized a triple-negative breast cancer patient-derived xenograft (PDX), PNX0204, from a patient that harbored a hemizygous germline BRCA1 mutation (1105insTC); the wild type BRCA1 allele was lost in the tumor, following a Loss of Heterozygosity model. PNX0204 tumors were originally sensitive to cisplatin treatment. After several rounds of cisplatin treatment and serial passage in mice, resistant tumors developed. Isogenic sensitive and resistant tumors were then tested for S1 sensitivity, with PEO1 (**Figure 3F**) and MDA-MB-436 (**Figure S2G**) xenografts serving as controls. As predicted, we found that the DNA fibers of cisplatin-sensitive PDX cells were degraded by S1 nuclease, but the fibers of cisplatin-resistant isogenic PDX cells were not, indicating ssDNA gaps had been suppressed in resistant patient samples (**Figure 3F**). Notably, resistant PDX suppressed gaps either by gap filling (**Figure 3F**), or by restored fork slowing (**Figure S2H**), indicating that loss of ssDNA gaps had occurred in BRCA patient tumors *de novo* and accurately predicted acquired cisplatin resistance (**Figure 3F and S2H**).

These findings present the idea that ssDNA gaps underlie BRCAness and chemosensitivity, and that loss of FP or HR do not. If so, when gaps are present, it should be possible to uncouple FP and HR from therapy response. To test this prediction, we first restored FP by inhibition of MRE11 or depletion of SMARCAL1 in BRCA2-deficient PEO1 cells (Kolinjivadi et al., 2017; Schlacher et al., 2011; Taglialatela et al., 2017). Nevertheless, even though FP was restored, cisplatin resistance was not conferred and, as predicted by our model, ssDNA gaps remained as demonstrated by the S1 nuclease degradation (**Figure 4A,B and S3A-E**). Moreover, neither SMARCAL1 nor MRE11 was predictive of BRCA2 cancer patient response based on mRNA levels in the TCGA database (**Figure 4C**) suggesting that gaps, but not FP, determine therapy response. Consistent with this point, cisplatin resistance as well as GS is re-established in BRCA2-mutant V-C8 cells upon complementation with either wild-type or the FP-defective C-terminal BRCA2 S3291A mutant (**Figure S3F,G**) (Schlacher et al., 2011). In addition to uncoupling FP from therapy response, these findings indicate that fork degradation occurs independent of replication gaps.

**Figure 4:**
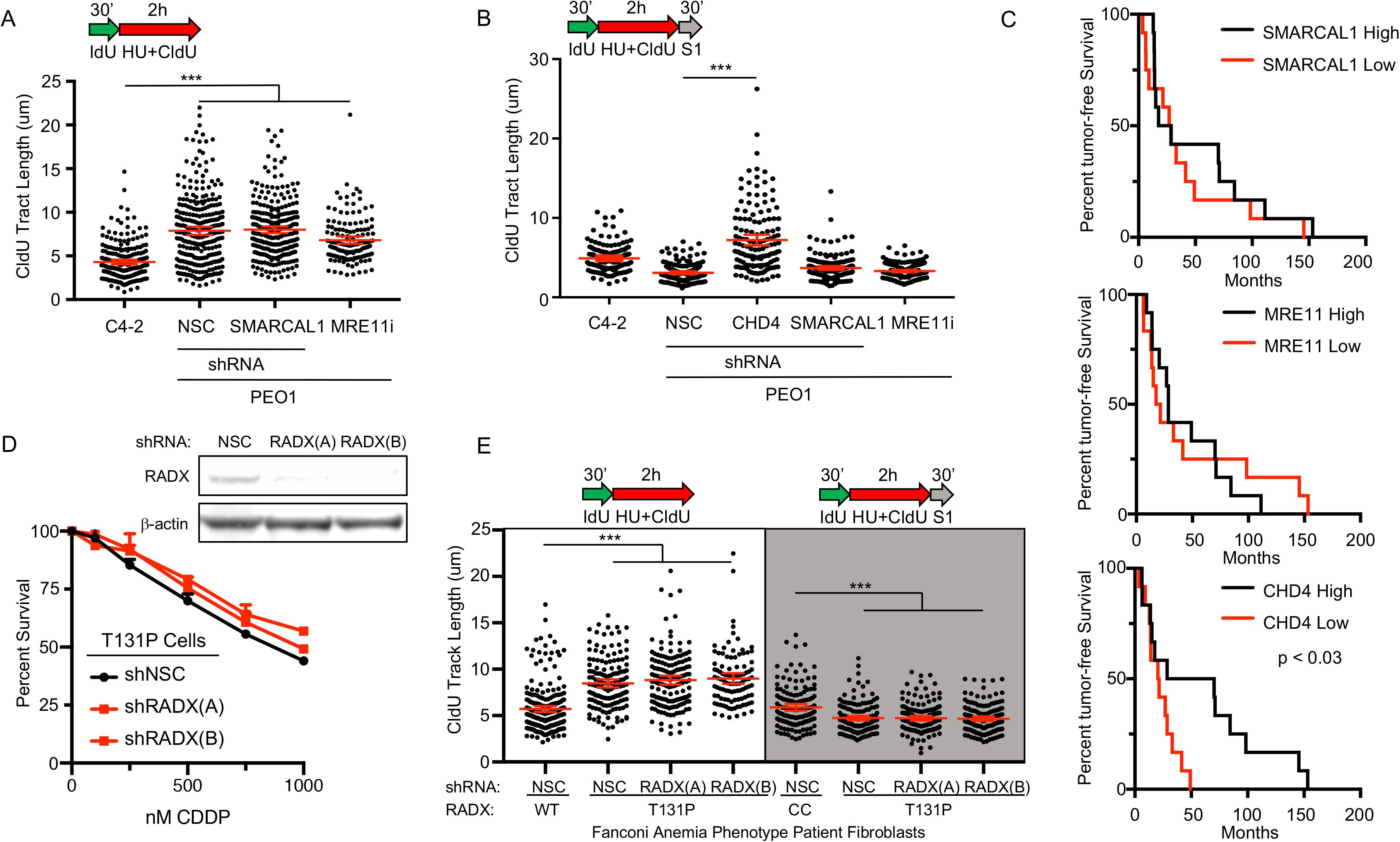
ssDNA replication gaps, and not FP or HR, determine patient response to chemotherapy and define BRCAness. **A)** Schematic and quantification of CldU track length in PEO1 cells shows that depleted SMARCAL1 or inhibited MRE11 does not increase replication in the presence of stress, and **B)** does not protect from S1 nuclease as found for CHD4 depletion. **C)** Neither SMARCAL1 nor MRE11 mRNA levels predict response of BRCA2 deficient ovarian cancer patients in TCGA dataset. In contrast, CHD4 mRNA levels do predict response in these patients. **D)** Top, Western blot confirms RADX is depleted by two shRNA reagents in T131P cells compared to non-silencing-control (NSC). Bottom, cell survival assay confirms RAD51 T131P cells remain hypersensitive to cisplatin even when RADX is depleted. **E)** Schematic and quantification of CldU track length shows (white panel) that fibroblasts from a Fanconi Anemia-like patient with a mutant allele of RAD51 (T131P; HR proficient cells, cisplatin sensitive) fail to arrest replication in the presence of stress even when RADX is depleted, and these regions are degraded by S1 nuclease (light grey panel). WT FA cells are corrected by CRISPR to delete the dominant-negative T131P RAD51 allele. Each dot represents one fiber. Experiments were performed in biological triplicate with at least 100 fibers per replicate; the xenograft fiber assay was performed in duplicate. Statistical analysis according to two-tailed Mann-Whitney test; *** p < 0.001. Mean and 95% confidence intervals are shown.

**Figure 5:**
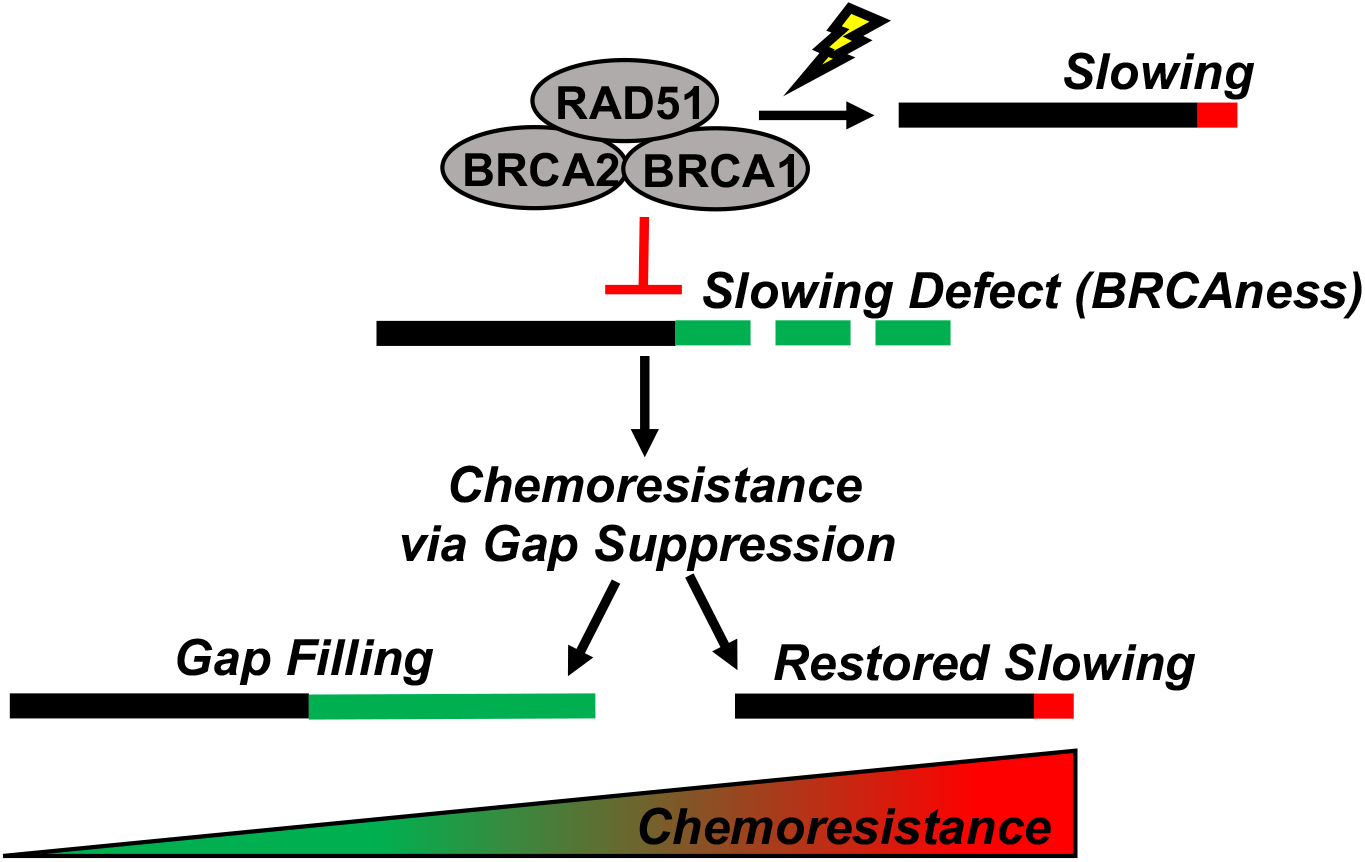
Model of BRCAness and chemosensitivity. During stress, BRCA-deficient cells fail to effectively restrain replication, leading to ssDNA gaps that determine chemosensitivity: BRCAness. These cells acquire chemoresistance by eliminating the ssDNA gaps, either by gap filling, or by restoring fork slowing.

We also considered the possibility that our ssDNA gap model could explain a discrepancy in the literature in which cells from a patient with Fanconi Anemia (FA) were sensitive to cisplatin and other genotoxic agents as expected, but were surprisingly found to be proficient in HR (Wang et al., 2015). Indeed, we found wide-spread ssDNA gap induction in the S1 assay in these FA patient cells; specifically, we observed S1 sensitivity in the FA patient fibroblasts that maintain a RAD51 mutant (T131P) allele as compared to isogenic RAD51 wild type fibroblasts (CRISPR corrected after isolation from the patient) (**Figure S4A and 4E**). However, because the FA cells are also deficient in FP, the sensitivity of the FA cells to cisplatin could be explained by either the induction of ssDNA gaps or loss of FP; indeed, wide-spread DSBs due to FP defects could in principle overwhelm HR. Thus, we restored FP in FA cells by depletion of the RAD51 negative regulator, RADX as previously reported (Bhat et al., 2018). However, despite restored FP and proficient HR, we nevertheless still observed cisplatin sensitivity and ssDNA gaps as before (**Figure 4D,E; S4B**), indicating that HR and FP cannot confer therapy resistance when gaps remain.

## Discussion

Our ssDNA model of chemoresponse has several implications. Namely, that replication gaps underlie the mechanism of action of chemotherapies, and it is the failure to suppress gaps, and not defects in HR or FP, that underlies the sensitivity of BRCA deficient cancer to therapy. In support of this concept, when gaps persist, we demonstrate that HR and/or FP proficient cells are nevertheless sensitive to cisplatin. Moreover, when gaps are suppressed by loss of CHD4, FEN1, or EZH2, HR deficient BRCA2 mutant cells are resistant to cisplatin (Guillemette et al., 2015; Meghani et al., 2018; Rondinelli et al., 2017), suggesting chemoresponse is independent of HR. Although gaps are a common indicator of replication stress and result from loss of the BRCA-RAD51 pathway, they have been overlooked as the determinant of toxicity in favor of defects in HR and FP (Hashimoto et al., 2010; Henry-Mowatt et al., 2003; Kolinjivadi et al., 2017; Sugimura et al., 2008; Wang et al., 2015; Xu et al., 2018; Zellweger et al., 2015). However, it is unclear how HR and FP participate in therapy response given that drugs used to treat BRCA deficient tumors do not initially cause DNA breaks or stall forks (Huang et al., 2013;Maya-Mendoza et al., 2018; Mutreja et al., 2018). In contrast, replication gaps arising due to loss of the BRCA-RAD51 pathway provides a logical explanation for the initiating lesion that is exacerbated by chemotherapies that further dysregulate replication.

In summary, this study indicates that cancer cells with deficiency in the BRCA pathway will be effectively treated by therapies that exacerbate replication gaps. Moreover, preventing gap suppression pathways will improve the effectiveness of chemotherapy as well potentially re-sensitize chemoresistant disease to therapy. Our findings also highlight that gaps could be biomarkers of BRCAness, and gap induction could be fundamental to the mechanism of action of chemotherapies that dysregulate replication.

## ACKNOWLEDGEMENTS

We thank the members of the Cantor laboratory for helpful discussions. We thank Dr. Agata Smogorzewska for providing the FA patient cells, Toshi Taniguchi for the PEO1 and C4-2 cells, Maria Jasin for the VC-8 cells, and Lee Zhou for the HCC1937 cells. This work was supported by R01 CA176166-01A1 (Cantor), R01 CA214799 and OC130212 (Johnson) as well as charitable contributions from Mr. and Mrs. Edward T. Vitone, Jr. and the Lipp Family Foundation. The Vermont Genetics Network Proteomics Facility is supported through NIH grant P20GM103449 from the INBRE Program of the National Institute of General Medical Science. We thank Drs. Igor Astutarov and Vladimir Khazak for establishing and sharing PNX0204.

## AUTHOR CONTRIBUTIONS

S.C., N.J. and N.P. designed the experiments. N.P., J.K., M.M., S.N., S.B., J.C., M.P., K.C., M.D., and M.B. performed the experiments. N.P., J.K., J.O., and L.J.Z. analyzed the data. S.C. and N.P. wrote the manuscript. S.C. and N.J. supervised the research.

## SUPPLEMENTAL FIGURE LEGENDS

**Figure S1:**
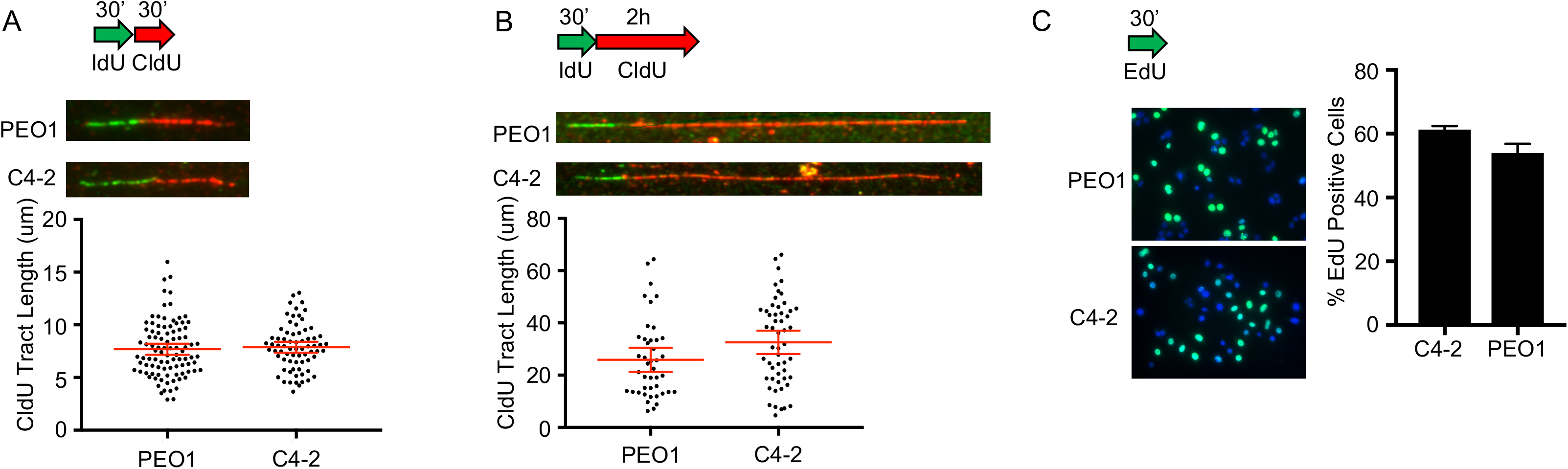
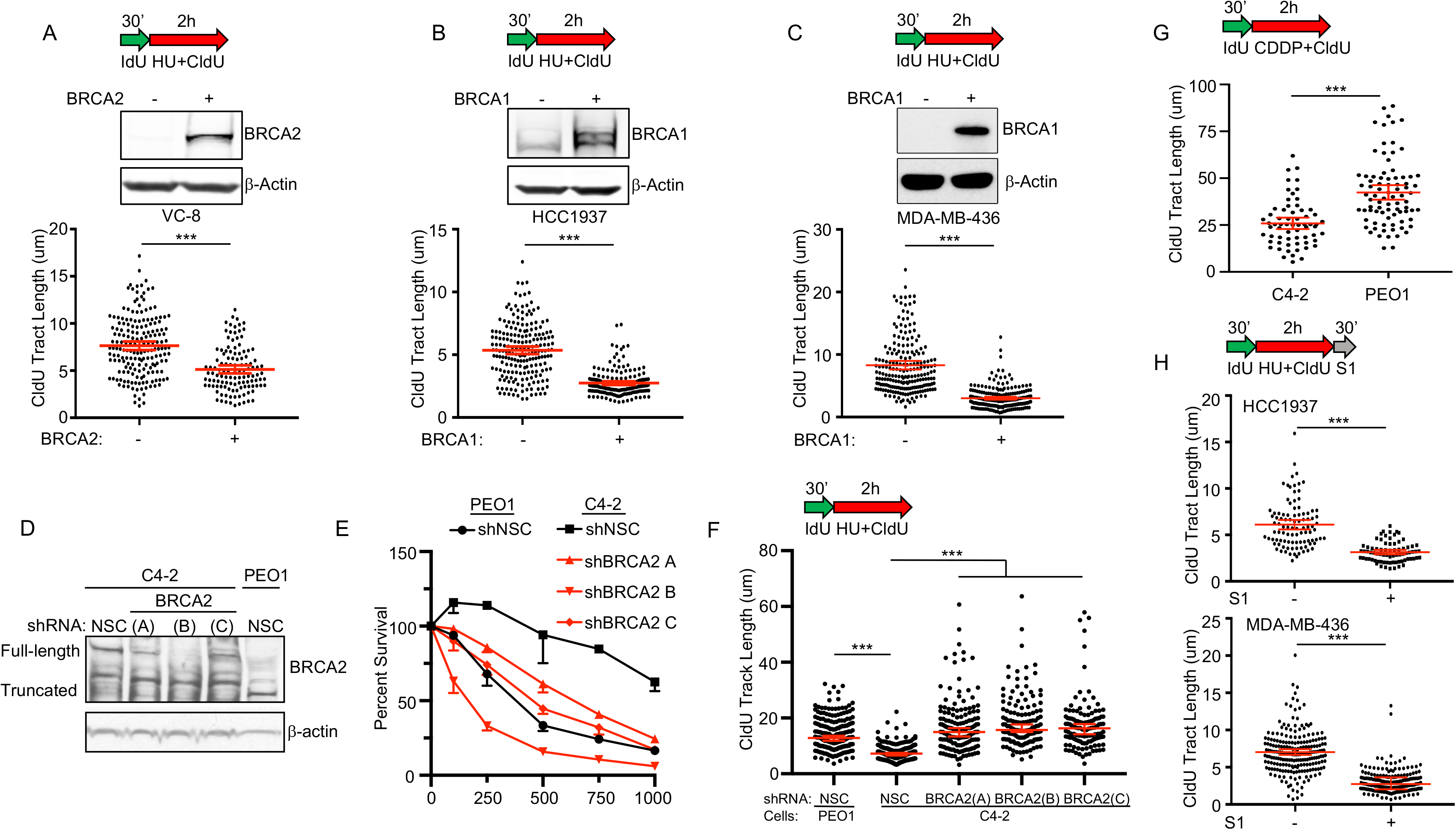
BRCA deficient cancer cells fail to restrain replication in the presence of stress and ssDNA gaps develop. **A)** Schematic, representative images, and quantification of CldU track length shows that PEO1 and C4-2 have similar track lengths in untreated conditions. **B)** Same as A, but with a 2h pulse. **C)** Schematic, representative images and quantification of nuclear imaging identifies a similar percentage of EdU positive cells in PEO1 as compared to C4-2 in unchallenged conditions. **S1.1A)** Western blot confirms BRCA deficiency and complementation in BRCA2 deficient VC-8 hamster cells and BRCA1 deficient breast cancer cells **S1.1B)** (HCC1937) and **S1.1C)** (MDA-MB-436). Schematic (above) and quantification of CldU track length shows BRCA complementation restores fork restraint. **S1.1D)** Western blot confirms BRCA2 is depleted in C4-2 by three distinct shRNA reagents. **S1.1E)** Cell survival assay confirms C4-2 cells with BRCA2 depleted are hypersensitive to cisplatin compared to C4-2-shNSC cells (with PEO1-shNSC as a hypersensitivity control). **S1.1F)** Schematic (above) and quantification of CldU track length shows BRCA2 depletion in BRCA2-proficient C4-2 creates a defect in slowing replication in response to HU as observed for BRCA2-deficient PEO1 cells. **S1.1G)** Schematic and quantification of CldU track length shows BRCA2 deficient PEO1 cells fail to effectively arrest replication in response to 1000nM CDDP compared to BRCA2 proficient C4-2 cells. **S1.1H)** Schematic and quantification of CldU track length shows BRCA1 deficient HCC1937 (top) and MDA-MB-436 (bottom) are sensitive to S1 nuclease. Each dot represents one fiber. Experiments were performed in biological duplicate with at least 100 fibers per replicate. Statistical analysis according to two-tailed Mann-Whitney test; *** p < 0.001. Mean and 95% confidence intervals are shown.

**Figure S2:**
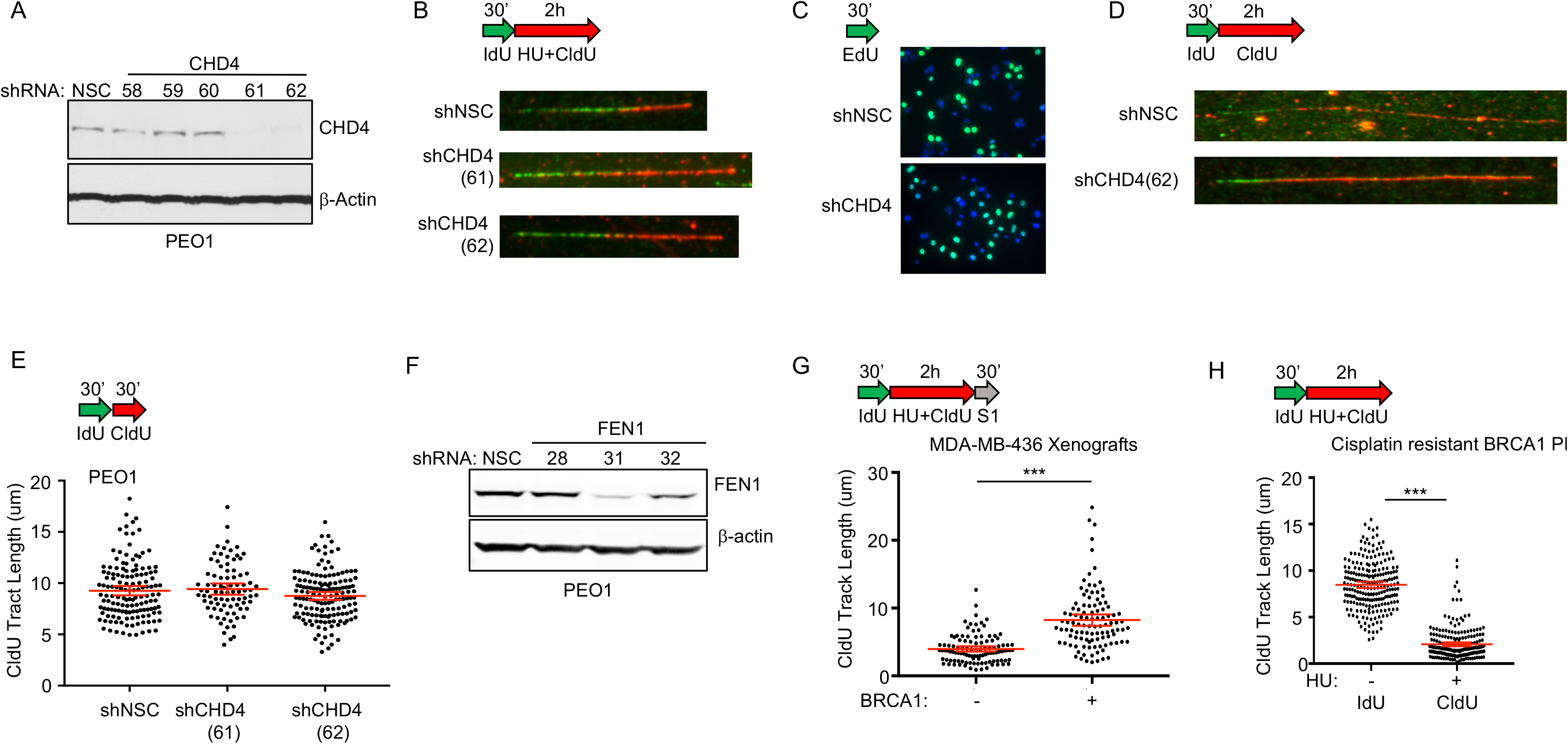
Replication restraint controls and depletion analysis. **A)** Western blot confirms CHD4 is depleted by shRNA (#61 and #62) compared to non-silencing control (NSC) in BRCA2 deficient PEO1. **B)** Schematic and representative images of CIdU track length shows that CHD4 depletion increases replication in the presence of stress. See Figure 2B and 4E for track lengths quantification for #62 and #61, respectively. **C)** Schematic and representative images of nuclear imaging show that CHD4 shRNA and NSC have similar numbers of EdU positive cells in unchallenged conditions. **D)** Schematic and representative images show that CHD4 shRNA and NSC have similar track lengths in unchallenged conditions with 2h labeling or **E)** 30 minute labeling. **F)** Western blot of shRNA confirms FEN1 is depleted by shRNA (#31 and #32) compared to non-silencing control (NSC) in BRCA2 deficient PEO1. **G)** Schematic and quantification of CIdU track length in MDA-MB-436 xenografts shows that complementation with BRCA1 protects from S1 nuclease. **H)** Schematic and quantification of DNA track length shows replication is efficiently arrested in BRCA1 deficient patient derived xenografts that have acquired chemoresistance. Each dot represents one fiber. Experiments were performed in biological duplicate with at least 100 fibers per replicate. Statistical analysis according to two-tailed Mann-Whitney test; *** p < 0.001. Mean and 95% confidence intervals are shown.

**Figure S3:**
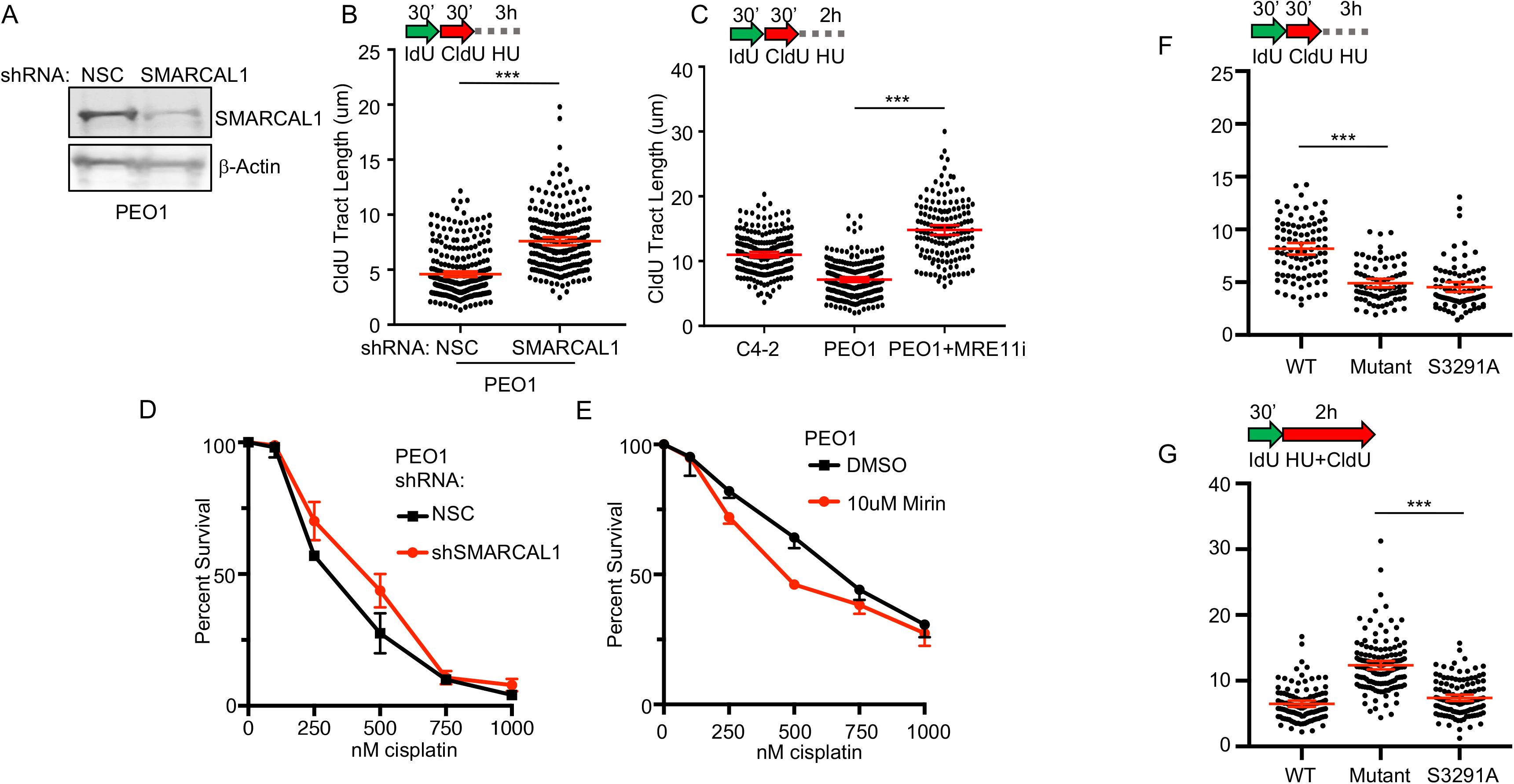
Depletion of SMARCAL1 or inhibition of MRE11 protects replication forks, but does not suppress ssDNA gaps or predict poor patient response. **A)** Western blot confirms SMARCAL1 is depleted in BRCA2-deficient PEO1 by shRNA as compared to NSC. **B, C)** Schematic and quantification of CldU track length shows that PEO1 with depleted SMARCAL1 or inhibited MRE11 protects replication forks after exposure to stress. **D)** cell survival assay confirms that SMARCAL1 depletion or **E)** MRE11i in PEO1 cells does not confer cisplatin resistance. **F)** Schematic and quantification of CldU track length shows that VC-8 complimented with BRCA2 S3291A are deficient for replication fork protection. **G)** Schematic and quantification of CldU track length shows that VC-8 complimented with BRCA2 S3291A are proficient for replication fork slowing. Each dot represents one fiber. Experiments were performed in biological duplicate with at least 100 fibers per replicate. Statistical analysis according to two-tailed Mann-Whitney test; *** p< 0.001. Mean and 95% confidence intervals are shown.

**Figure S4:**
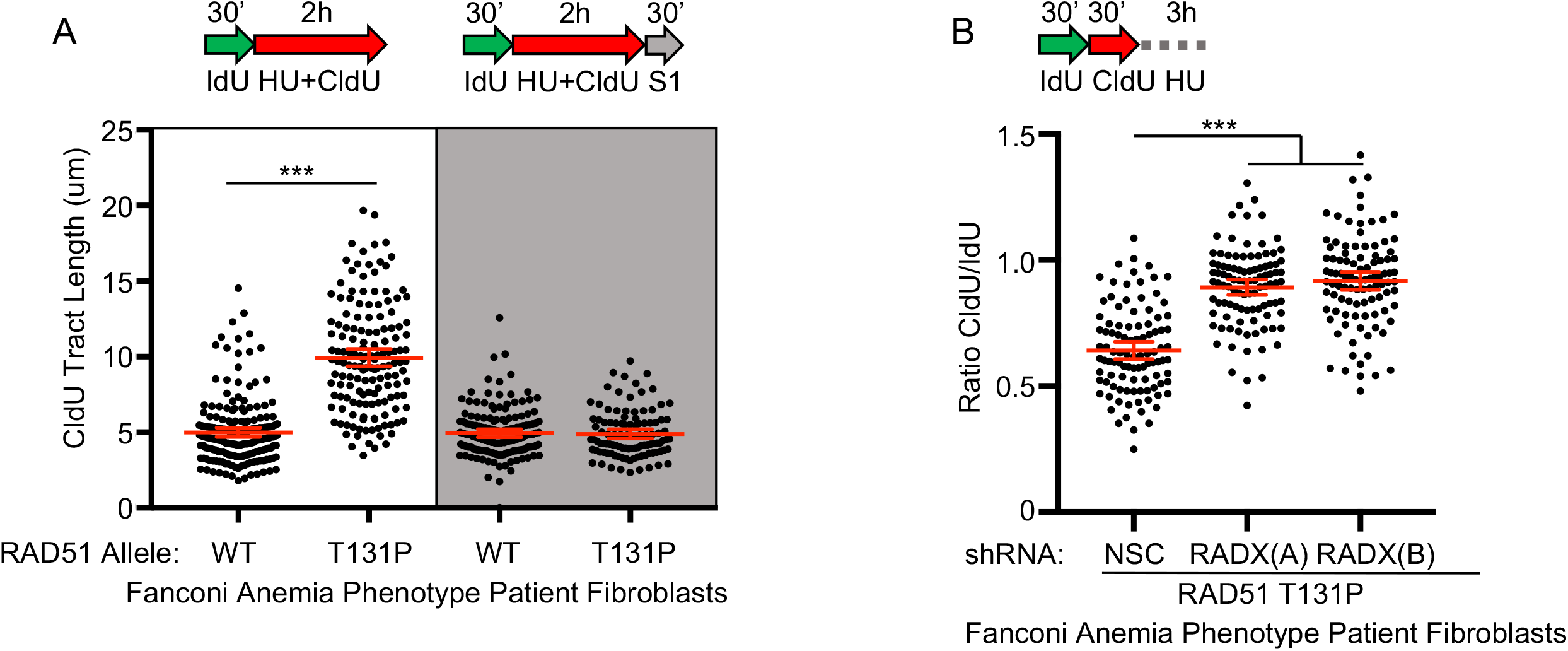
Fanconi Anemia Patient Fibroblasts with a RAD51 T131P Mutant Allele (HR Proficient) Generate ssDNA Gaps even when Fork Protection is Restored. **A)** Schematic and quantification of CldU track length shows (white panel) that RAD51 T131P cells fail to arrest replication in the presence of stress compared to the CRISPR corrected control that deletes the dominant T131P allele. These regions are degraded by S1 nuclease (light grey panel). **B)** Schematic and quantification of CldU track length shows that RAD51 T131P cells are FP deficient, and depletion of RADX confers FP. Each dot represents one fiber. Experiments were performed in biological triplicate with at least 100 fibers per replicate. Statistical analysis according to two-tailed Mann-Whitney test; *** p< 0.001. Mean and 95% confidence intervals are shown.

## MATERIALS AND METHODS

### Cell Culture

PEO1, C4-2, VC-8, and MDA-MB-436 cell lines were cultured in DMEM + 10% FBS + 1% P/S. HCC1937 Deficient and HCC1937 + WT BRCA1 were cultured in RPMI1640 + L-Glutamine + 10% FBS + 1% P/S.

### Drugs

Cisplatin was prepared as a 1mM solution in saline per the manufacturer’s instructions (Sigma). HU was prepared fresh in complete media prior to experiments per the manufacturer’s instructions. The MRE11 inhibitor mirin (Sigma) was prepared as a 50mM solution in DMSO and used at 50uM for DNA fiber analysis and was added during the indicated step per the figure diagrams (Schlacher et al., 2011). EZH2 was inhibited with 5uM GSK126 (Selleck) and was added during the CldU + HU step in S1 and slowing experiments (Rondinelli et al., 2017). 5-chloro-2’-deoxyuridine (CldU) and 5-lodo-2’-deoxyuridine (IdU) were obtained from Sigma.

### shRNA

HEK293T cells were used to package lentiviral particles with the pLKO.1 shRNA system as previously described(Guillemette et al., 2015). Briefly, HEK293 cells were transfected with 1:1:2 μg of packaging plasmids versus shRNA hairpins on the pLKO.1 vector using Effectene transfection reagent (Qiagen) 48 h prior to harvesting supernatants. Supernatants were filtered and added to recipient cell lines with 1 μg/mL polybrene. Cells infected with shRNA vectors were selected with puromycin. For shRNA-mediated silencing, the following hairpins from The RNAi Consortium were obtained from GE Dharmacon:

CHD4-61, TRCN0000021361: 5’-GCTGACACAGTTATTATCTAT-3’

CHD4-62, TRCN0000021362: 5’-GCTGACACAGTTATTATCTAT-3’

ZFHX3-58, TRCN0000013558: 5’-GCCAGGAAGAATTATGAGAAT-3’

FEN1-32, TRCN0000049732: 5’-GATGCCTCTATGAGCATTTAT-3’

SMARCAL1-69, TRCN0000083569: 5’-GCGGAACTCATTGCAGTGTTT-3’

### CellTiter-Glo 2.0 Toxicity Assays

Cells were plated at 500 cells per well in the center wells of a 96 well plate in 200ul volume and allowed to adhere for 36h. Subsequently, drugs were added in a 100ul volume and the cells were incubated for 10 days. CellTiter-Glo 2.0 (Promega) was used to quantify cell number by ATP. To prevent evaporation, all blank wells in the 96 well plate were filled with media, and each plate was placed in a humid chamber with PBS.

### DNA Fiber Assays

DNA fiber assays were performed as previously described. Briefly, cells were plated at 10^6 cells per 10cm dish and allowed to adhere for 36h. Subsequently, DNA was labeled for 30 minutes with 50uM IdU and washed with PBS, and treated with 50uM CldU and replication stress depending on the assay. For fork restraint assays, cells were exposed to 50uM CldU with 0.5mM HU for 2h. For fork restraint with continued stress, cells were exposed to 50uM CldU with 0.5mM HU for 2h, followed by 4mM HU for 2-3h. For fork degradation assays, cells were labeled with 50uM CldU alone, followed by 4mM HU for 3-5h. After labeling, cells were collected with trypsin, washed with PBS, and resuspended in PBS at 250,000 cells/ml. 2ul of cell solution was placed on a positively charged slide, followed by lysis for 8 minutes with 12.5ul of spreading buffer (0.5% SDS, 200mM Tris-HCl, pH 7.4, 50mM EDTA). Slides were tilted to a 45 degree angle to allow fibers to spread, allowed to dry for 20 minutes, fixed in 3:1 Methanol:Acetic Acid for 3 minutes, rehydrated in PBS for 5 minutes, denatured with 2.5mM HCl for 30 minutes, blocked with PBS + 0.1% TritonX-100 + 3% BSA for 1h, and treated with primary (2.5h, 1:100) and secondary antibodies (1h, 1:200) in PBS + 0.1% TritonX-100 + 3% BSA. Track lengths were measured in Fiji (Schindelin et al., 2012). The antibody used to detect IdU was anti-BrdU (Becton Dickinson 347580, detects both BrdU and IdU); the antibody used to detect CldU was anti-BrdU (Abcam ab6326, detects both BrdU and CldU). For the nondenaturing fiber assay to detect ssDNA, all steps are the same, except acid steps were removed (both acetic acid from the fixation step, and the HCl denaturing step). In addition, IdU was replaced with EdU and detected by ClickIT EdU Alexa 488 Imaging Kit (Thermo Scientific) to label analog in non-denatured DNA per the manufacturer’s instructions. For total nuclear imaging, cells 100,000 cells were plated onto poly-l-lysine coated coverslips and allowed to adhere. Cells were subsequently treated with EdU as indicated, fixed and processed for Click-IT EdU detection according to the manufacturer’s instructions (Thermo Scientific). Coverslips were mounted in Vectashield with DAPI, and ten fields were imaged at 20x in the center of the coverslip. EdU intensity per cell was quantified with Cell Profiler from the Broad Institute(Carpenter et al., 2006).

### S1 Nuclease Fiber Assay

As described previously, cells were exposed to 50uM IdU to label replication forks, followed by 50uM CldU with 0.5mM HU for 2h. Subsequently, cells were permeabilized with CSK buffer (100 mM NaCl, 10 mM MOPS, 3 mM MgCl2 pH 7.2, 300 mM sucrose, 0.5% Triton X-100) at room temperature for 8 minutes, followed by S1 nuclease (20U/ml) in S1 buffer (30 mM Sodium Acetate pH 4.6, 10 mM Zinc Acetate, 5% Glycerol, 50 mM NaCl) for 30 minutes at 37C. Finally, cells were collected by scraping, pelleted, resuspended in 100-500ul PBS; 2ul of cell suspension was spotted on a positively charged slide and lysed and processed as described in the DNA fiber assay section above.

### Cell Fractionation

To isolate cytoplasmic, nuclear, and chromatin fractions for western blot or mass spectrometry, cells were lysed with the NE-PER kit according to the manufacturer’s instructions (Thermo Scientific). To isolate chromatin fractions, the insoluble pellet that remains from NE-PER lysis was resuspended in 60ul of 2x loading buffer with DTT, heated at 70C for 10 minutes, and sonicated in a BioRuptor (Diagenode) for 20 minutes on high, with a cycle of 30 seconds on and 30 seconds off.

### Western Blot

All steps were performed according to the manufacturer’s instructions (Thermo Scientific). The protein concentration of different cellular fractions was determined by BCA Assay. Samples were reduced with DTT in LDS loading buffer, and heated at 70C for 10 minutes. 40ug total protein was fully resolved on either a Tris-Acetate gel (for large proteins) or a Bis-Tris gel (for small proteins), and transferred to a nitrocellulose membrane. The nitrocellulose membrane was processed for near-infrared quantitative westerns according to the manufacturer’s protocol (LiCor). The membrane was allowed to dry for 20 minutes, total protein was stained with the REVERT stain as a total protein loading control, followed by blocking with Odyssey blocking buffer, treated with primary antibody overnight, followed by near-infrared secondary antibody (800CW) at 1:5000 for 1h. The membrane was allowed to dry prior to imaging on the LiCor Odyssey Imager. Primary antibodies used include anti-BRCA2 (Abcam ab123491, 1:1000); anti-CHD4 (Abcam ab54603 1:1000); anti-CHD4 (Abcam ab70469, for immunoprecipitation); anti-SMARCAL1 (Abam ab37003, 1:1000); anti-ZFHX3 (Lifespan Biosciences LS-C179898-100); anti-PCNA (Abcam ab29); anti-B-actin (Sigma A5441, 1:15,000).

### Proteomics

For SILAC, PEO1 were dual labeled in SILAC media with dialyzed FBS (Thermo Scientific) with heavy lysine (K+8) and heavy arginine (R+10) from Cambridge Isotope Labs. PEO1 and C4-2 (unlabeled SILAC media) were treated with cisplatin, collected with trypsin, counted, mixed at a 1:1 ratio, and fractionated together in the same Eppendorf tube with the NE-PER kit as described. Cellular fractions were fully resolved on SDS-PAGE gels, fixed with Imperial Protein Coomassie Stain (Thermo Scientific), washed in water overnight, and cut into 13 molecular weight regions corresponding to the protein marker standard. Each region was reduced with DTT, alkylated with iodoacetamide, and digested with Trypsin Gold for Mass Spectrometry with ProteaseMAX according to the manufacturer’s instructions (Promega). Peptides were dried in a speedvac, resuspended in 6 ul buffer A (0.1% formic acid), and 2 ul tryptic digests were analyzed on the Thermo Q-Exactive mass spectrometer coupled to an EASY-nLC Ultra system (Thermo Fisher). Peptides were separated on reversed phase columns (12 cm x 100 μm I.D), packed with Halo C18 (2.7 um particle size, 90 nm pore size, Michrom Bioresources) at a flow rate of 300 nl/min with a gradient of 0 to 40% acetonitrile (0.1% FA) over 55 min. Peptides were injected into the mass spectrometer via a nanospray ionization source at a spray voltage of 2.2 kV. The mass spectrometer was operated in a data-dependent fashion using a top-10 mode(Peng et al., 2018).

### Processing of Proteomics Data

Raw proteomics data were analyzed with MaxQuant software (Cox and Mann, 2008). We required a false discovery rate (FDR) of 0.01 for proteins and peptides and a minimum peptide length of 7 amino acids. MS/MS spectra were searched against the human proteome from UniProt. For the Andromeda search, we selected trypsin allowing for cleavage N-terminal to proline as the enzyme specificity. We selected cysteine carbamidomethylation as a fixed modification, and protein N-terminal acetylation and methionine oxidation were selected as variable modifications. Two missed tryptic cleavages were allowed. Initial mass deviation of precursor ion was up to 7 ppm, mass deviation for fragment ions was 0.5 Dalton. Protein identification required one unique peptide to the protein group. Known contaminants were removed from the analysis. To identify statistical significance, isotopic ratios of identified proteins from three biological replicates were analyzed using the *limma* statistical package (Ritchie et al., 2015). The isotopic ratio obtained from MaxQuant was subsequently converted to log2 scale and plotted against the −log10(p-value) for each gene in GraphPad Prism.

### TCGA Database Analysis

The TCGA database was used to identify ovarian cancer patients with germline mutations in BRCA2, and subsequently tested for correlation between patient progression free survival and the mRNA expression of genes of interest. To obtain patient germline sequencing data, we applied for access to protected TCGA patient data through NIH. The germline BAM sequencing data for each patient at the BRCA2 locus was downloaded. The BAM Slicing tool option and the GDC API were used to automate the process. Sliced BAM files were sorted and indexed using SAMTOOLS (Li et al., 2009), and mutations were identified using the Genome Analysis Tool Kit (GATK), Broad Institute (McKenna et al., 2010). We followed the GATK best practices for germline mutation calling until the last step of the protocol, where we used hard filtering instead of variant quality score recalibration (VQSR) because sliced BAM files at a single locus are not compatible with VQSR. Briefly, we called germline variants using the HaplotypeCaller tool in GVCF mode, consolidated the GVCF files using the GenomicsDBImport Tool, and called mutations using the GenotypeGVCFs tool. The data were hard filtered to isolate BRCA2 germline mutations, and we classified mutations using the Variant Effect Predictor (VEP) from Ensembl (McLaren et al., 2016). Finally, mutations were selected that are predicted to disable BRCA2, including premature stop codons, frameshift mutations, and deletions. A case list of patient barcodes was compiled harboring at least one of these BRCA2 disabling mutations, and cBioPortal (Cerami et al., 2012; Gao et al., 2013) was used to obtain the progression free survival data and mRNA expression data for our genes of interest. Patients with mRNA expression of the target gene over the median were classified as high expression; patients below were classified as low expression. The survival curve of the low and high expression groups was plotted in GraphPad Prism, and significance was determined using the Log-rank test.

### PDX Methods

PNX0204 was derived at Fox Chase Cancer Center under IRB and IACUC approved protocols. PDX tumors were grown in *NOD.Cg-Prkdc^scid^ Il2rg^tm1Wjl^/SzJ* (NSG) mice. Cisplatin resistant PDX tumors were obtained from mice after tumors progressed on serial treatments of 6 mg/kg cisplatin. The tumors were harvested at approximately 500 mm^3^ and dissociated in 0.2% collagenase, 0.33 mg/ml dispase solution for 3h at 37°C. The dissociated cells were maintained at 37°C in RPMI1640 + 10% FBS and used for DNA fiber assays within 24h of tumor extrackion. DNA fiber and S1 nuclease fiber assays were performed as described above.

